# Human Surfactant Protein D Binds S1 and Receptor Binding Domain of Spike protein and acts as an entry inhibitor of SARS-CoV-2 Pseudotyped viral particles *in vitro*

**DOI:** 10.1101/2020.12.18.423418

**Authors:** Miao-Hsi Hsieh, Nazar beirag, Valarmathy Murugaiah, Yu-Chi Chou, Wen-Shuo Kuo, Hui-Fan Kao, Taruna Madan, Uday Kishore, Jiu-Yao Wang

## Abstract

Human SP-D is a potent innate immune molecule whose presence at pulmonary mucosal surfaces allows immune surveillance role against pulmonary pathogens. Higher levels of serum SP-D have been reported in patients with severe acute respiratory syndrome coronavirus-1 (SARS-CoV). Studies have suggested the ability of human SP-D to recognise spike glycoprotein of SARS-CoV; its interaction with HCoV-229E strain leads to viral inhibition in human bronchial epithelial (16HBE) cells. Previous studies have reported that a recombinant fragment of human SP-D (rfhSP-D) composed of 8 Gly-X-Y repeats, neck and CRD region, can act against a range of viral pathogens including influenza A Virus and Respiratory Syncytial Virus *in vitro, in vivo* and *ex vivo* models. In this context, this study was aimed at examining the likely protective role of rfhSP-D against SARS-CoV-2 infection. rfhSP-D showed a dose-responsive binding to S1 spike protein of SARS-CoV-2 and its receptor binding domain. Importantly, rfhSP-D inhibited interaction of S1 protein with the HEK293T cells overexpressing Angiotensin Converting Enzyme 2. The protective role of rfhSP-D against SARS-CoV-2 infection as an entry inhibitor was further validated by the use of pseudotyped lentiviral particles expressing SARS-CoV-2 S1 protein; ~0.5 RLU fold reduction in viral entry was seen following rfhSP-D treatment (10 μg/ml). The results highlight the therapeutic potential of rfhSP-D in SARS-CoV-2 infection and merits pre-clinical studies in murine models.

## Introduction

Human surfactant protein D (SP-D), a collagen-containing C-type lectin and a member of collectin family, which is known to be involved in surfactant homeostasis and pulmonary immunity (1). SP-D is primarily synthesized and secreted into the air space of the lungs by alveolar type II and Clara cells (2, 3). Its primary structure is organized into four regions: a cysteine-rich N-terminus, a triple-helical collagen region composed of Gly-X-Y triplets repeats, an α-helical coiled neck region, and a C-terminal C-type lectin or carbohydrate recognition domain (CRD) (1). As a versatile innate immune molecule, SP-D can interact with a number of pathogens triggering clearance mechanisms against viruses, bacteria, and fungi, as well as and apoptotic cells (4).

Direct interaction of SP-D with a wide range of viruses results in viral neutralization, and induction of phagocytosis *in vitro* (5) (6). Anti-viral activity of human SP-D against Influenza A Virus (IAV) infection has been reported. SP-D binds haemagglutinin (HA) and neuraminidase (NA) glycoproteins of IAV, and thus, inhibits hemagglutination at initial stages of the infection (7). A recombinant fragment of human SP-D (rfhSP-D), composed of homotrimeric neck and three CRD region, was also shown to bind HA, NA and Matrix 1 (M1) viral proteins of IAV, and act as an entry inhibitor of IAV infection on A549 lung epithelial cells (8). Furthermore, SP-D binds to gp120 and inhibits HIV-1 infectivity and replication (9) in U937 monocytic cells, Jurkat T cells and PBMCs, in addition to suppressing HIV-1 triggered cytokine storm (10). Higher levels of serum SP-D have been detected in patients infected with severe acute respiratory syndrome (SARS) coronavirus-1 (SARS-CoV) (11). SARS-CoV spike glycoprotein is recognized by SP-D (12). Interaction between SP-D and HCoV-229E, another coronavirus strain, leads to inhibition of viral infection in human bronchial epithelial (16HBE) cells (13).

SARS coronavirus 2 (SARS-CoV-2) is an enveloped β-coronavirus, belonging to the Coronaviridae family of viruses, and is genetically close to SARS-CoV (~80% sequence similarity) and bat coronavirus RaTG13 (96.2%) (14, 15). The envelope of SARS-CoV-2 is coated by the spike (S) glycoprotein, small envelope (E) glycoprotein, membrane (M) glycoprotein, nucleocapsid (N) protein, and several putative accessory proteins (15–17). The SARS-CoV-2 mediates its entry into the host cell using the S1 sub-unit of the S glycoprotein by binding to angiotensin-converting enzyme 2 (ACE2) receptor (18). However, viral entry into the host cells requires not only binding to the ACE2 receptor, but also priming of the S protein by a transmembrane protease serine 2 (TMPRSS2) via cleavage of the S protein at S1/S2 sites (19). This cleavage process is very crucial for the virus-host cell membrane fusion and cell entry (20). Following viral replication, assembly, and release, the infected host cells undergo pyroptosis, thus, releasing Damage-Associated Molecular Patterns (DAMPs) (21). DAMPs are then recognised by surrounding macrophages and monocytes that respond to viral infection by inducing cytokine storm (22). However, in some cases, an impaired or dysregulated immune response can also occur, causing an Acute Respiratory Distress Syndrome (ARDS) (23, 24).

Designing and developing new anti-viral therapeutic strategies are crucial for treating SARS-CoV-2. The likely anti-viral effects of immune-surveillance molecules like rfhSP-D have not been well investigated in SARS-CoV-2 infection. Since rfhSP-D has been shown to inhibit viral infection and replication of IAV and HIV-1, this study was aimed at investigating whether rfhSP-D can interfere with the binding of SARS-CoV-2 S1, and receptor binding domain (RBD) of SARS-CoV-2 with ACE-2. The ability of rfhSP-D to inhibit infection and replication of SARS-CoV-2 was also examined using pseudotyped lentiviral particles expressing SARS-CoV-2 S1 protein.

## Material and methods

### Expression and Purification of rfhSP-D

DNA sequences coding for 8 Gly-X-Y repeats of collagen region, α-helical neck and CRD region of human SP-D were cloned under T7 promoter and expressed in *Escherichia coli* BL21 (λDE3) pLysS using construct pUK-D1 (25, 26). Primary bacterial inoculum (25 ml) was grown in Luria-Bertani (LB) medium (500 ml) with 34 μg/ml chloramphenicol and 100 μg/ml ampicillin (Sigma-Aldrich) at 37°C until an OD_600_ of 0.6 was reached. Following isopropyl β-D-thiogalactoside (IPTG) (0.5mM) induction, the transformed *E. coli* cells were grown further for another 3 h at 37 °C on a shaker. The bacterial cells were harvested by centrifugation (5000 rpm, 4°C, 10 min), and the cell pellet was resuspended in lysis buffer containing 50 mM Tris-HCl, pH 7.5, 200 mM NaCl, 5 mM EDTA, 0.1% v/v Triton X-100, 0.1 mM phenylmethane sulfonyl fluoride (PMSF) (Sigma-Aldrich), and 50 μg lysozyme (Sigma-Aldrich) at 4°C for 1 h. The lysed cell lysate was then sonicated at 60 Hz for 30 sec with an interval of 2 min (12 cycles) using a Soniprep 150 (MSE, London, UK), followed by centrifugation (12,000 rpm, 15 min). The inclusion bodies were denatured using buffer (50 ml) containing 0.5 M Tris-HCl, 0.1 M NaCl, pH7.5 and 8 M urea for 1 h at 4°C. The soluble fraction was dialysed against the same buffer containing varied concentration of urea (4 M, 2 M, 1 M, 0 M) for 2 h each. The refolded material was then dialysed against affinity buffer (50 mM Tris–HCl, pH7.5, 100 mM NaCl, 10 mM CaCl_2_) for 2 h at 4°C. The affinity buffer dialysed supernatant was then loaded on to a maltose-agarose column (5 ml) (Sigma-Aldrich); the bound rfhSP-D was eluted using elution buffer containing 50 mM Tris-HCl, 100 mM NaCl, 10 mM EDTA. Purified rfhSP-D was run on SDS-PAGE to assess its purity. LPS was removed using Endotoxin Removal Resin (Sigma-Aldrich). LPS level was determined using QCL-1000 Limulus amebocyte lysate system (Lonza) and found to be >5 pg/ μg of rfhSP-D.

### SDS-PAGE

To detect the purity of purified rfhSP-D fractions, an SDS-PAGE (12% v/v) gel was used. Purified rfhSP-D was diluted in 1:1 (v/v) ratio in Laemmli sample buffer [1 M Tris-HCl (pH 6.8, 1 ml), 10% w/v SDS (4 ml; Sigma-Aldrich), 98 % v/v Glycerol (2 ml; Fisher-Scientific), 1% w/v Bromophenol blue, β-mercaptoethanol (2.5 ml; Sigma-Aldrich) and distilled water (d.H_2_O). The protein sample was then denatured at 95°C for 10 min before loading on to the gel. To assess the size of purified rfhSP-D, a standard pre-stained protein marker (Fisher Scientific) was also loaded. The SDS-PAGE gel was stained for 2 h using staining solution [Brilliant Blue (1 g; Sigma-Aldrich), methanol (50% v/v; Fisher-Scientific), acetic acid (10% v/v; Fisher-Scientific) and D.H_2_O (40 ml). The stained gel was then de-stained using the de-staining solution [methanol (40% v/v), acetic acid (10% v/v; Fisher-Scientific) until the protein bands were visible.

### Western Blotting

To determine the immunoreactivity of purified rfhSP-D, purified rfhSP-D (10 μg) was resuspended in Laemmli sample buffer (10 μl) and heated at 100°C for 10 min. Bovine serum albumin (BSA; Thermo-Fisher) was used as a negative control protein. The heated samples were subjected to 12% (v/v) SDS-PAGE and then electrophoretically transferred onto a nitrocellulose membrane (320mA for 2h) (Sigma-Aldrich) in transfer buffer [25mM Tris–HCl pH 7.5, 190 mM glycine (Sigma-Aldrich), and 20% v/v methanol (Fisher-Scientific)]. The membrane was incubated in 5% w/v dried milk powder (Sigma-Aldrich) diluted in PBS for 2h at 4°C to block non-specific binding. After blocking, the membrane was washed with PBS three times (5 mins each wash). The membrane was incubated with polyclonal rabbit anti-human SP-D primary antibody (1:1000; MRC Immunochemistry Unit, Oxford) for 1h at room temperature. The unbound primary antibody was washed off using PBS+ 0.05% v/v Tween 20 (PBST) (3 times, 10 min each wash). The membrane was then probed with secondary Goat anti-rabbit IgG horseradish peroxidase (HRP)-conjugate (1:1000; Fisher Scientific) for 1h at room temperature. Following PBST washes, the membrane-bound rfhSP-D was visualized by developing the membrane using 3’-diaminobenzidine (DAB) substrate kit (Thermo-Scientific).

### ELISA

Polystyrene microtiter plates (Sigma-Aldrich) were coated with SARS-CoV-2 spike S1 protein (NativeAntigen) or RBD (Acro) (5 μg/ml) at 4°C overnight using carbonate/bicarbonate (CBC) buffer, pH 9.6 (Sigma-Aldrich). The following day, the microtitre wells were washed three times with Tris Buffered Saline-Tween (TBST, pH 7.2-7.4) containing 0.05% v/v Tween 20 (Sigma-Aldrich) and 5mM CaCl_2_ (Thermo-Scientific),. The wells were then blocked by TBS containing 1% w/v BSA and 5mM CaCl_2_, for 1 h. After washing three times with TBST, the wells were incubated with two-fold dilutions of rfhSP-D protein in the blocking buffer at 4°C overnight. Next day, the wells were washed and then incubated with human SP-D detecting antibody (0.5 μg/ml) (R&D Systems; 1:180) for 2 h at room temperature. After washing, the wells were incubated with streptavidin-HRP (1:40; R&D Systems) for 20 min, followed by washing three times. TMB substrate (100 μl/well; Thermo-Fisher) was added to each well and the reaction was stopped using 1M H_2_SO_4_ (50μl/well; Sigma-Aldrich). In a parallel experiment, the wells were washed and then incubated with polyclonal rabbit anti-human SP-D primary antibody (0.5 μg/ml) (1:5000) for 2 h at room temperature. After washing, the wells were incubated with Goat anti-rabbit IgG-HRP-conjugate (1:5000; Fisher Scientific) for 1 h, followed by washing three times. TMB substrate (100 μl/well; Thermo-Fisher) was added to each well and the reaction was stopped using 1M H_2_SO_4_ (100 μl/well; Sigma-Aldrich). Absorbance at 450nm were measured by VersaMax™ ELISA Microplate Reader.

### Competitive ELISA

Polystyrene microtiter plates were coated with 2 μg/ml rfhSP-D at 4°C overnight using CBC buffer and washed three times with TBS buffer containing 0.05% v/v Tween 20 and 5mM CaCl_2_. The wells were blocked with TBS containing 1% BSA and 5mM CaCl_2_ for 1h. The wells were then washed three times and incubated with SARS-CoV-2 spike S1 protein (sheep-IgG tag) or RBD (His-tag) (2.5 or 5 μg/ml) separately in blocking buffer containing 10mM maltose and 10mM EDTA at 4°C overnight. Next day, the wells were washed and then incubated with anti-sheep IgG-HRP antibodies or anti-His antibodies (Genetex, GTX628914, 0.5 μg/ml) (1:2000) for 2 h. For the detection of RBD binding, the wells were further incubated with anti-mouse IgG antibody (Abcam, ab6728, 0.5 μg/ml) (1:2000) for 2 h. After washing, the plates were incubated with TMB substrate (100 μl/well) and then quenched with 1M H_2_SO_4_ (50 μl/well). Absorbance at 450nm was recorded by VersaMax™ ELISA Microplate Reader.

### Cell culture and treatments

Human embryonic kidney (HEK) 293T or HEK293T cells overexpressing ACE2 receptor (HEK293T-ACE2) were cultured in complete Gibco Dulbecco’s Modified Eagle Medium (DMEM), supplemented with 10% v/v fetal bovine serum (FBS), 100 U/ml penicillin (Sigma-Aldrich) and 100 μg/ml streptomycin (Sigma-Aldrich), and left to grow at 37°C in the presence of 5% v/v CO_2_ for approximately 48 h before passaging. Since HEK293T cells were adherent, they were detached using 2× Trypsin-EDTA (0.5%) (Thermo Fisher Scientific) for 10 min at 37°C. Cells were then centrifuged at 1,500 rpm for 5 min, followed by re-suspension in complete DMEM medium. To determine the cell count and viability, an equal volume of the cell suspension and Trypan Blue (0.4% w/v) (Thermo Fisher Scientific) solution were vortexed, followed by cell count using a hemocytometer with Neubauer rulings (Sigma-Aldrich). Cells were then re-suspended in complete DMEM for further use.

### Flow cytometry

ACE2 expression was assessed between HEK293T cells overexpressing ACE2 receptor (HEK293T-ACE2) and HEK293T cells using flow cytometry. Briefly, both ACE2-transfected and non-transfected HEK293T cells (1×10^5^ cells) were incubated with ACE2 antibody [N1N2], N-term (GeneTex, GT×101395) (1:250) for 1 h at room temperature.

Following PBS washes, the cells were probed with goat anti-Rabbit IgG (H+L) Cross-Adsorbed Secondary Antibody linked to Alexa Fluor 647 (Thermo Fisher Scientific) (0.6 μl/100 μl per tube) for 1 h at room temperature in dark. After washing the cells with PBS, the cells were resuspended in FACS buffer (PBS containing 2% FBS) and subjected to flow cytometry.

For binding experiments using rfhSP-D, SARS-CoV-2 S1 protein containing a C-terminal His-tag (Acro; S1N-C52H3) (5 μg/ml) was tagged with anti-His antibody (Genetex; GT395) (1:100) at 4°C for 1h and followed by pre-incubation with a series of two-fold dilutions of rfhSP-D (10 μg/ml) or mock (cells only) at 4°C for 1h. HEK293T-ACE2 cells (1×10^5^ cells) were incubated in DMEM incomplete medium with the mixture of SARS-CoV-2 S1 protein, anti-His antibodies and rfhSP-D at 37°C for 2 h. The cells were collected and washed with FACS buffer twice and incubated with anti-mouse IgG-PE (Genetex, GTX25881) conjugate (1:100) for 30 min and washed three times. The live cells were gated from FSC vs. SSC dot plot in order to determine the PE expressed cells containing S1 on their surface by CytoFLEX.

### Fluorescence Microscopy

HEK293T and HEK293T-ACE2 cells (0.5 × 10^5^) were grown on coverslips in complete DMEM medium overnight under standard culture conditions, as mentioned above. Next day, cells were washed with PBS three times, the coverslips were fixed with 4% v/v paraformaldehyde (Sigma-Aldrich) for 15 minutes, and then washed twice. The coverslips were permeabilized with 0.25% v/v Triton-100 (Sigma-Aldrich) for 15 min. After washing, coverslips were blocked with 2% w/v BSA for 1h and incubated with ACE2 antibody [SN0754] (1:250) (GeneTex, GTX01160), followed by Goat anti-rabbit IgG (H+L) cross-adsorbed secondary antibody (1:500) (Thermo Fisher Scientific) for 1 h at room temperature in dark. After incubation with secondary antibody, the cells were washed twice with PBS and mounted in the medium with DAPI (Abcam) on the slides to visualize under an upright fluorescence microscope (BX51; Olympus).

### Production of SARS-CoV-2 Pseudotyped Lentivirus

The pseudotyped lentivirus carrying SARS-CoV-2 spike protein was generated by transiently transfecting HEK293T cells with pCMV-DR8.91, pLAS2w.Fluc.Ppuro and pcDNA3.1-nCoV-SD18 (SARS-CoV-2 spike gene with 54 nucleotides deletion at its C-terminus was synthesized and cloned into pcDNA3.1 expression vector). HEK293T cells were seeded one day before, and then transfected with the indicated plasmids using TransIT®-LT1 transfection reagent (Mirus). The culture medium was replenished at 16 h and harvested at 48h and 72h post-transfection. Cell debris was removed by centrifugation at 4,000 x g for 10 min, and the supernatant was passed through 0.45-mm syringe filter (Pall Corporation). The pseudotyped lentivirus was aliquoted and stored at −80°C until further use. The transduction unit (TU) of SARS-CoV-2 pseudotyped lentivirus was estimated using cell viability assay in response to the limited dilution of lentivirus. In brief, HEK293T cells, stably expressing human ACE2, were plated on 96-well plate one day before lentivirus transduction. For titrating, different amounts of lentivirus particles were added to the culture medium containing polybrene (final concentration 8 mg/ml). Spin infection was carried out at 1,100 x g in 96-well plate for 30 min at 37°C. After Incubating cells at 37°C for 16 h, the culture medium containing virus particles and polybrene was removed and replaced with fresh complete DMEM containing 2.5 μg/ml puromycin. After treating with puromycin for 48 h, the culture media was removed, and the cell viability was assessed using 10% AlarmaBlue reagents, according to manufacturer’s instruction. The survival of uninfected cells (without puromycin treatment) was set as 100%. The virus particle titer (transduction units) was determined by plotting the survival of cells versus diluted viral dose.

### Pseudotyped virus neutralization assay

HEK293T cells in 10 cm dishes were transfected with pCMVΔR8.91, pcDNA nCoV-SD18 and pLAS2w.FLuc.Ppuro plasmids (5, 2, 8 μg, respectively). Next day, cells were washed with PBS gently, and replaced with 10 ml of fresh medium (RPMI containing 10% FBS). The medium at 48 and 72 h were collected and stored in −80°C for future use. HEK293T-ACE2 cells (HEK293T cells overexpressing ACE2 receptor) (0.5×10^5^ cells) were preincubated with rfhSP-D (0, 5, 10 and 20 μg/ml) for 24 h and then washed twice with PBS. The SARS-CoV-2 pseudotyped lentiviral particle containing medium (500μl/well) was added on to the cells, followed by incubation at 37°C under standard culture conditions. After 2h, fresh complete DMEM medium (500 μl) was added onto the cells and incubated at 37°C. Following 72 h incubation, the cells were washed with PBS twice, and incubated with lysis buffer at 37°C for 10 min. Firefly luciferase activity (RLU) was measured using ONE-Glo™ Luciferase Assay System (Promega) and FlexStation.

### Statistical Analysis

GraphPad Prism 6.0 software was used to generate all the graphs. Two-way ANOVA test was used for the statistical analysis. The significance values were considered between rfhSP-D treated and untreated conditions, based on *p<0.05. Error bars show the SD or SEM (figure legends).

## Results

### Expression, purification, and characterization of a recombinant fragment of human SP-D (rfhSP-D)

A recombinant truncated form of human SP-D, containing 8 Gly-X-Y triplets, α-helical coiled-coil neck domain, and CRD (rfhSP-D), was expressed under bacteriophage T7 promoter in *Escherichia coli* BL21(λ DE3) pLysS as inclusion bodies. Following IPTG induction, rfhSP-D was overexpressed at 20 kDa protein, and the inclusion bodies containing insoluble rfhSP-D were refolded via denaturation and renaturation process. The soluble rfhSP-D fractions were further purified using an affinity chromatography using a maltose agarose column, and then eluted with 10 mM EDTA, confirming that the correctly folded rfhSP-D bound maltose in a calcium-dependent manner(27) Under reducing conditions, rfhSP-D migrated as a monomer of **~**20 kDa on a 12% SDS-PAGE (v/v) (Figure 1A). The immunoreactivity of purified rfhSP-D protein was verified via western blotting analysis using rabbit polyclonal anti-human SP-D primary antibody that was raised against native human SP-D purified from lung lavage of alveolar proteinosis patients (Figure 1B).

**Figure 1:**
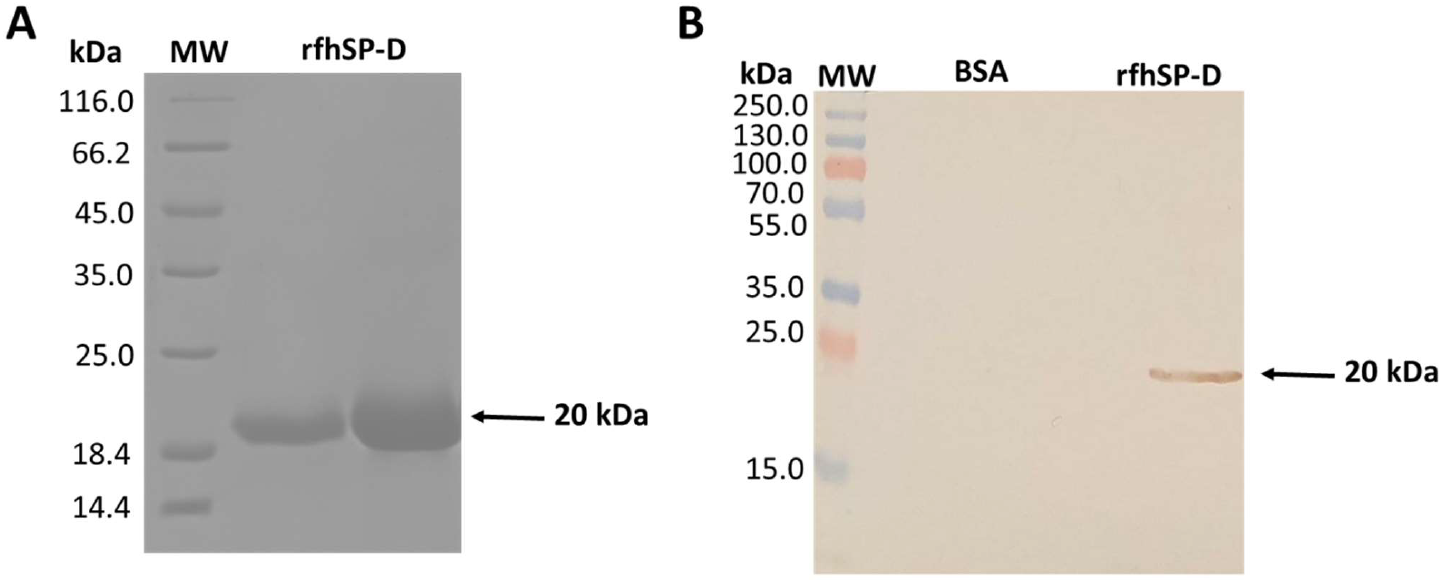
12% SDS-PAGE (v/v) (A) and immunoblot profile (B) of affinity purified LPS-free recombinant form of human surfactant protein D (rfhSP-D) under reducing condition. (A) pUK-D1 composed of the neck and CRD DNA sequence was expressed under the bacteriophage T7 promoter, which expressed rfhSP-D as ~20 kDa insoluble protein. rfhSP-D inclusion bodies were subjected to denaturation and renaturation cycle using a gradient of urea buffer and the dialysate was passed through a maltose–agarose column. The eluted protein appeared as a single band at ~20 kDa. (B) The immunoreactivity of purified rfhSP-D protein was verified via western blotting analysis using rabbit polyclonal anti-human SP-D primary antibody (1:1000) that was raised against native human SP-D purified from lung lavage of alveolar proteinosis patients; lane 1: BSA as a negative control; lane 2: purified LPS-free rfhSP-D (5 μg/well).

### Interaction of rfhSP-D with S protein of SARS-CoV-2

In SARS-CoV, S-protein is the predominant surface glycoprotein recognized by the host innate immune mechanisms. The S protein of SARS-CoV-2 has almost 76% identity to SARS-CoV. Previous studies indicated that SP-D bound S protein of SARS-CoV-1 which required Ca^2+^; the binding was inhibited by maltose. Therefore, the first part of the study was aimed at examining the interaction of rfhSP-D with spike protein (S1) using direct binding ELISA. It was found that rfhSP-D bound SARS-CoV-2 S1 protein in a dose-dependent manner; this interaction was inhibited by maltose and EDTA (Figure 2A). Among varied concentrations of rfhSP-D tested, a strong and maximum binding of rfhSP-D with SARS-CoV-2 S1 (5 μg/ml) was observed at 10 μg/ml.

**Figure 2:**
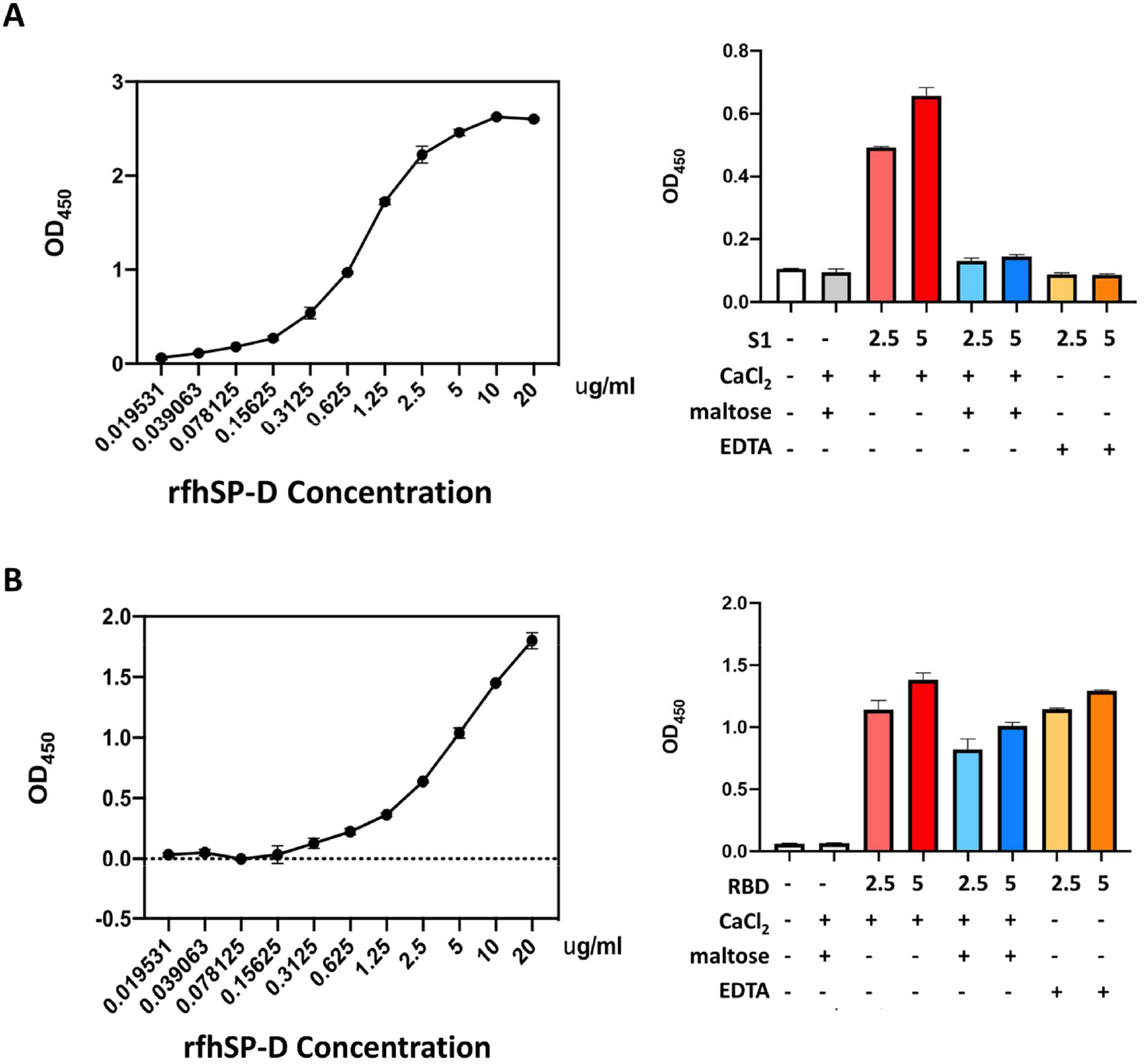
rfhSP-D binding with the spike (S1) (A) and its RBD of the SARS-CoV-2 (B) was determined via direct ELISA. Microtiter wells were coated with SARS-CoV-2 spike S1 protein (5 μg/ml) or RBD (5 μg/ml) in carbonate-bicarbonate buffer, pH 9.6 overnight at 4°C. The following day, the wells were blocked with Tris Buffered Saline (TBS) buffer containing 1% BSA and 5mM CaCl_2_, pH 7.2-7.4. After washing the wells with TBS, the wells were incubated with a series of two-fold dilutions of rfhSP-D protein in blocking buffer at 4°C overnight. The binding between S1 protein and rfhSP-D was detected using polyclonal anti-human SP-D antibody (1:5000), followed by probing with Goat anti-rabbit IgG horseradish peroxidase (HRP)-conjugate (1:5000). The data were expressed as mean of three independent experiments done in triplicates ± SEM.

### Interaction of rfhSP-D with the RBD of S protein of SARS-CoV-2

The binding of SARS-CoV-2 to its cellular receptor, ACE2, is mediated by the RBD region of the S protein. A higher binding affinity has been reported for RBD of SARS-COV-2 to ACE2 receptor compared to SARS-CoV (28). Furthermore, RBD of SARS-CoV-2 has been suggested to have a crucial role in spike protein-induced viral attachment, fusion, and entry into the host cells (29). In this context, this study was also aimed at determining the ability of rfhSP-D to bind RBD of SARS-CoV-2 (Figure 2B) via direct ELISA. rfhSP-D bound RBD in a dose-dependent manner, but the binding interaction was stronger compared to S1 protein. Furthermore, addition of calcium or chelation of Ca^2+^ by EDTA did not significantly affect the interaction between rfhSP-D and RBD region (Figure 3). However, the addition of maltose reduced binding efficiency (Figure 3). No rfhSP-D binding was observed in the absence of RBD, indicating a lack of non-specific interaction in this assay. These results suggest that the protein-protein interaction may occur between the CRD region of rfhSP-D and the RBD region of S protein in a calcium-independent manner.

**Figure 3:**
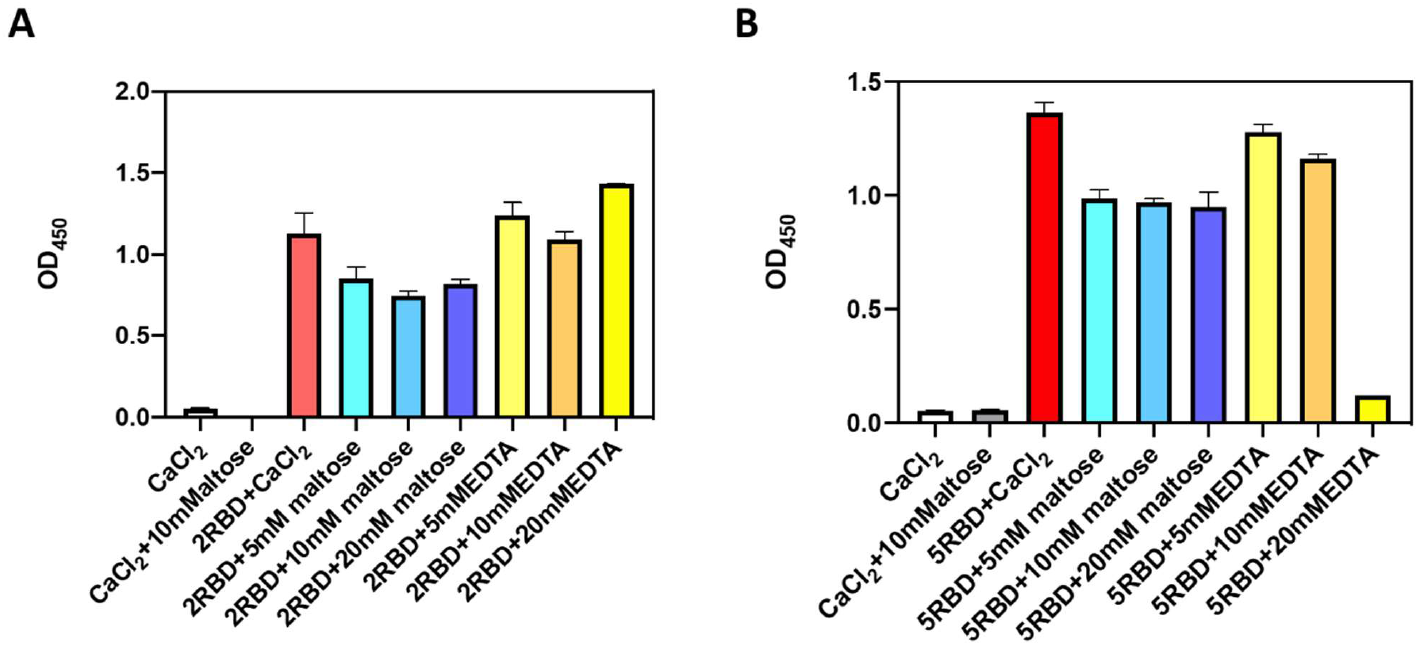
Competitive ELISA to show that rfhSP-D binding to S1 (A) and its RBD (B) of SARS-COV-2 interaction is inhibited by maltose. Polystyrene microtiter plates were coated with 2μg/ml rfhSP-D, and incubated with SARS-CoV-2 spike S1 protein (5 μg/ml) (sheep-IgG tag) or RBD (His-tag) (5 μg/ml). The binding was detected using antisheep IgG HRP antibodies (1:5000) or anti-His antibodies ((1:5000). Absorbance at 450nm were recorded by VersaMax™ ELISA Microplate Reader.

### rfhSP-D interaction with SARS-CoV-2 S1 and ACE2 overexpressing HEK293T cells

The S1 spike protein of the SARS-CoV-2 contains RBD that can recognise and interact with its cellular receptor, angiotensin-converting enzyme 2 (ACE2) (30, 31), thus mediating viral entry into the host cells. Since rfhSP-D was found to interact with the spike protein and its RBD at a protein level, we also tested the ability of rfhSP-D to interact with HEK293T cells overexpressing ACE2 receptor. Successful transfection of the ACE2 receptor gene into HEK293T cells was verified by measuring the expression levels of ACE2 receptor via immunofluorescence microscopy and flow cytometry (Figure 4). Quantitative and qualitative analysis of the ACE2 receptor using ACE2 antibody (SN0754) revealed a higher signal for ACE2 on HEK293T-ACE2 cells when compared to HEK293T cells alone. This study also focused on examining whether rfhSP-D treatment can inhibit the interaction of SARS-CoV-2 S1 with ACE2 receptor on HEK293T cells (Figure 5). Pre-incubation of SARS-CoV-2 S1 protein (2 μg/ml) with a varied concentration of rfhSP-D (0.625 −10 μg/ml), was found to reduce S1 binding of HEK293T cells overexpressing ACE2 receptor in a dose-dependent manner. The rfhSP-D treatment at 10 μg/ml was found to reduce the binding of S1 to ACE2 receptor on HEK293T cells by ~35% when compared to the control (S1+0 μg/ml rfhSP-D).

**Figure 4:**
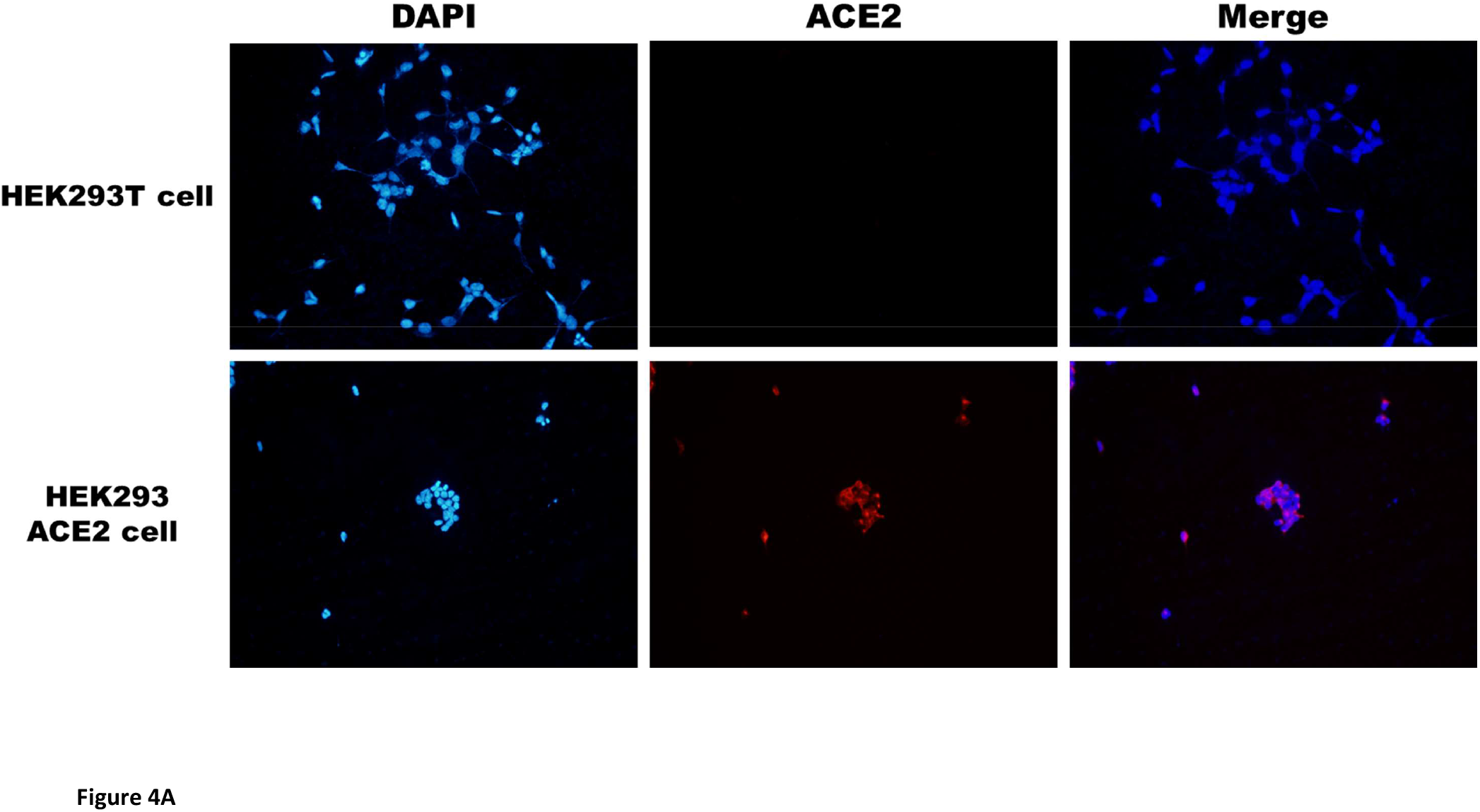
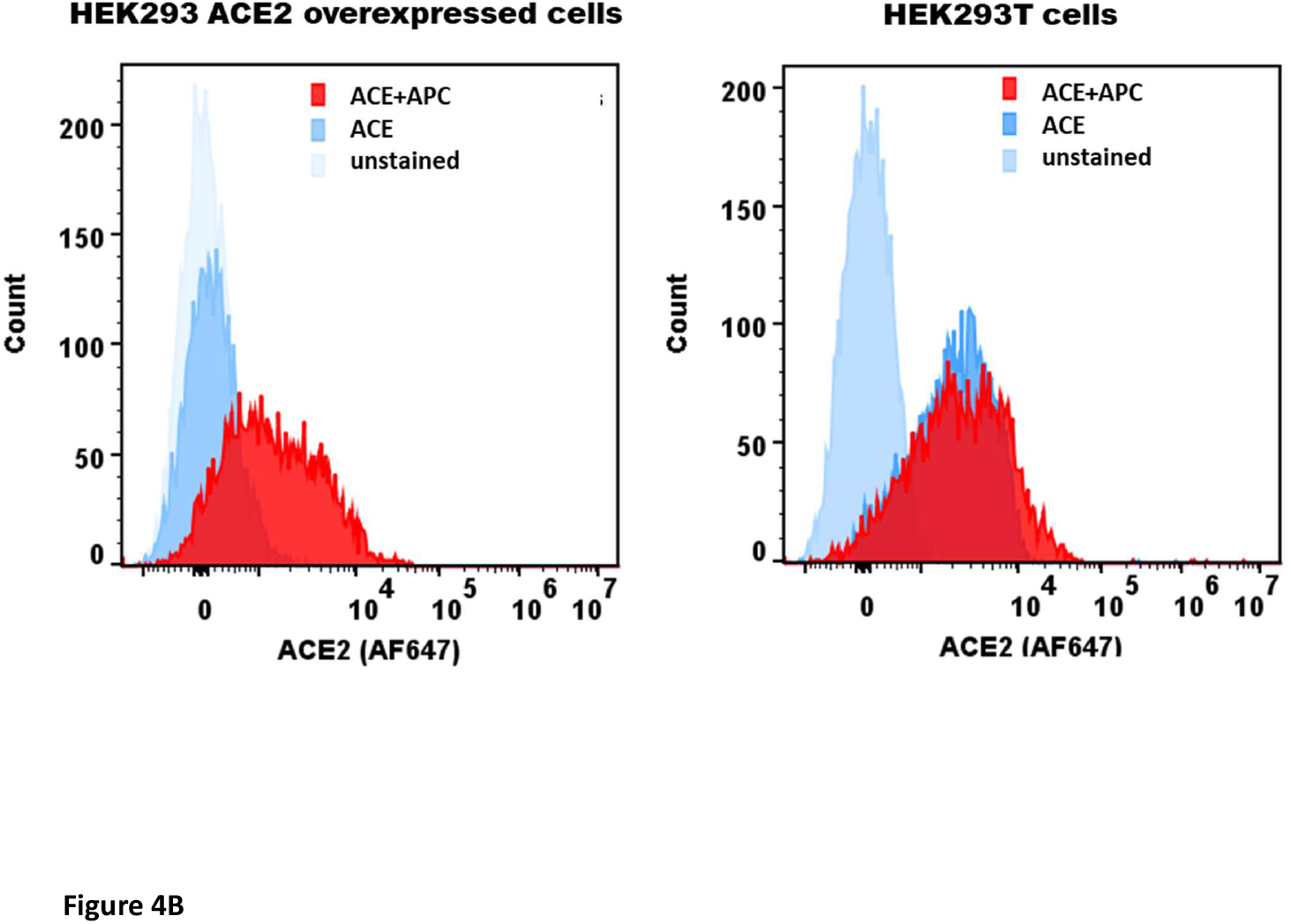
Expression of ACE2 receptor was determined via immunofluorescence microscopy (A) and flow cytometry (B). (A) HEK293T (0.5×10^5^ cells) and HEK293T-ACE2 cells (0.5×10^5^ cells) were seeded on coverslips, followed by incubation at 37°C under standard culture conditions. After wasing the cells with PBS twice, the ACE2 expression was detected in both cell lines using the ACE2 antibody [SN0754](1:250), followed byincubation for 1 h at room temperature. Following PBS washes, Goat antirabbit IgG (H+L) cross-adsorbed secondary antibody (1:500) was added. Following PBS washes, the coverslips were mounted in medium with DAPI on a microscopy slide and viewed under a fluorescence microscope (Olympus). (B) Flow cytometric analysis of ACE2 expression was determined by the the shift in the fluorescence intensity using ACE2 antibody [N1N2], N-term (GeneTex) (1:250). The ACE2 expression was detetected by CytoFLEX.

**Figure 5:**
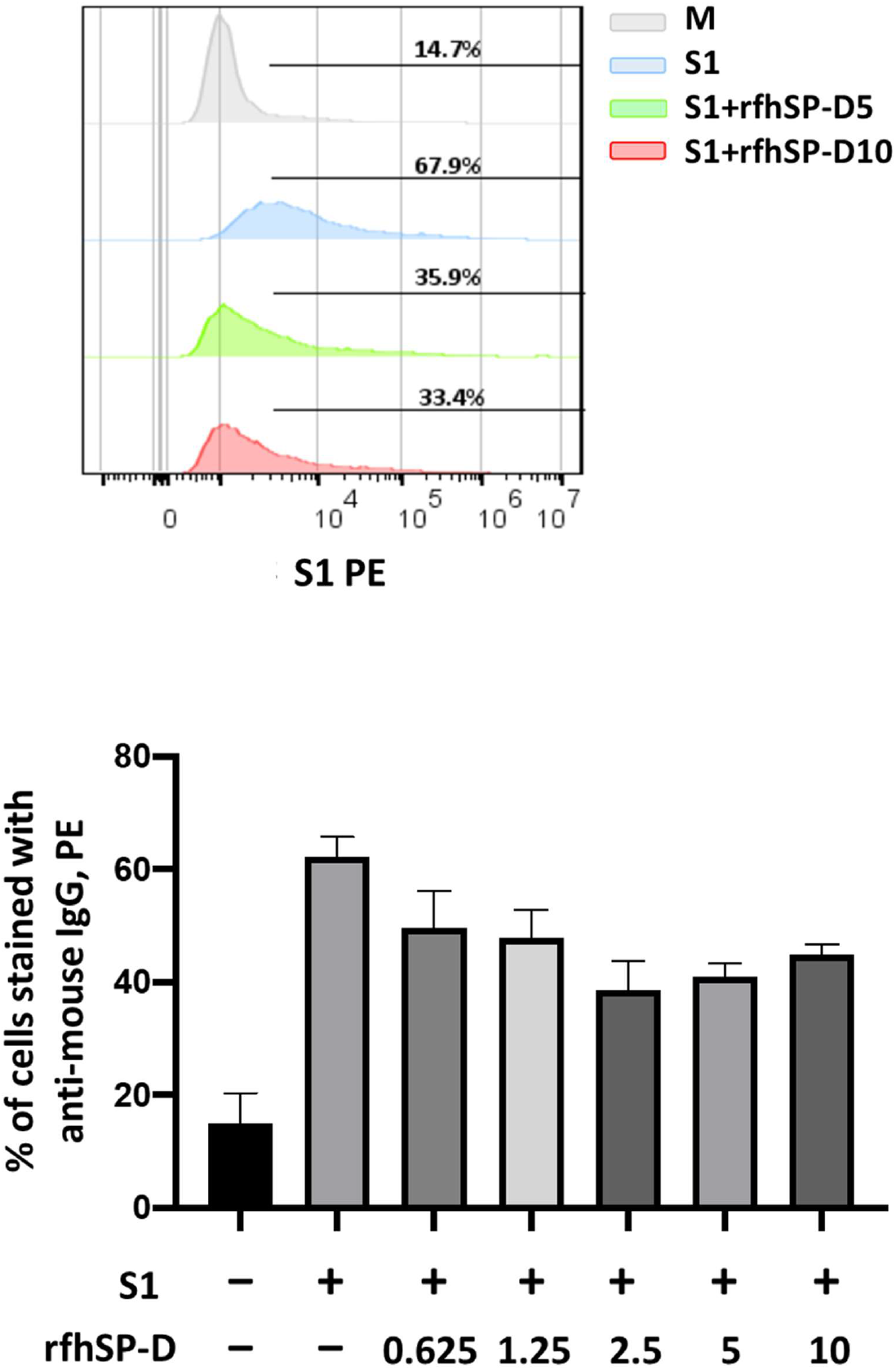
rfhSP-D treatment inhibits the interaction between SARS-CoV-2 S1 and ACE2 receptor on HEK293T cells. Protein complex was made by tagging SARS-CoV-2 S1 protein (5 ug/ml) with anti-His antibody (10ug/ml), followed by incubation with rfhSP-D (2.5, 5 or 10 μg/ml) for 2h at room temperature. This complex (S1+ anti-His+rfhSP-D) was added on to HEK293T-ACE2 cells (1×10^5^ cells) at 37 °C for 2 h. The cells were collected and washed with FACS buffer twice and incubated with anti-mouse IgGPE conjugate (Genetex, GTX25881) (1:100) for 30 min and washed three times. The cells stained with S1 were detected by CytoFLEX.

### rfhSP-D act as an entry inhibitor of SARS-COV-2 infection

After confirming the ability of rfhSP-D to prevent the interaction between SARS-CoV-2 S1 protein and HEK293T cells overexpressing ACE2 receptor, we investigated whether rfhSP-D modulated viral entry using a luciferase reporter assay using Pseudotyped lentiviral particles expressing SARS-CoV-2 S1 protein (Figure 6). SARS-CoV-2 pseudotyped lentiviral particles were produced as a safe strategy to study the involvement of S1 glycoprotein in the recognition and neutralization of the virus by a varied concentration of rfhSP-D. The production of lentiviral particles pseudotyped with envelope protein S1 was carried out by co-transfecting HEK293T cells with plasmid containing the coding sequence of the indicated pcDNA3.1-nCoV-SD18 (SARS-CoV-2 spike gene), pLAS2w.Fluc.Ppuro, and pCMV-DR8.91. Purified pseudotyped particles and cell lysate harvested at 48 and 72 h were analyzed via western blotting, and the expression level of SARS-CoV-2 spike protein was determined using anti-S1 monoclonal antibody (data not shown). Cells pre-incubated rfhSP-D (5 and 10 μg/ml) showed a significant ~ 0.5 RLU fold reduction in luciferase activity (1.0 × 10^5^ RLU) compared to the cells+SARS-CoV-2 (1.5 × 10^5^ RLU). The reduced luciferase activity following treatment with rfhSP-D indicated that the interaction between rfhSP-D and SARS-CoV-2 S1 protein interfered with S1-containing viral particle binding to ACE2, and hence, prevented the entry of the virus into the HEK323T-ACE2 cells (Figure 6).

**Figure 6:**
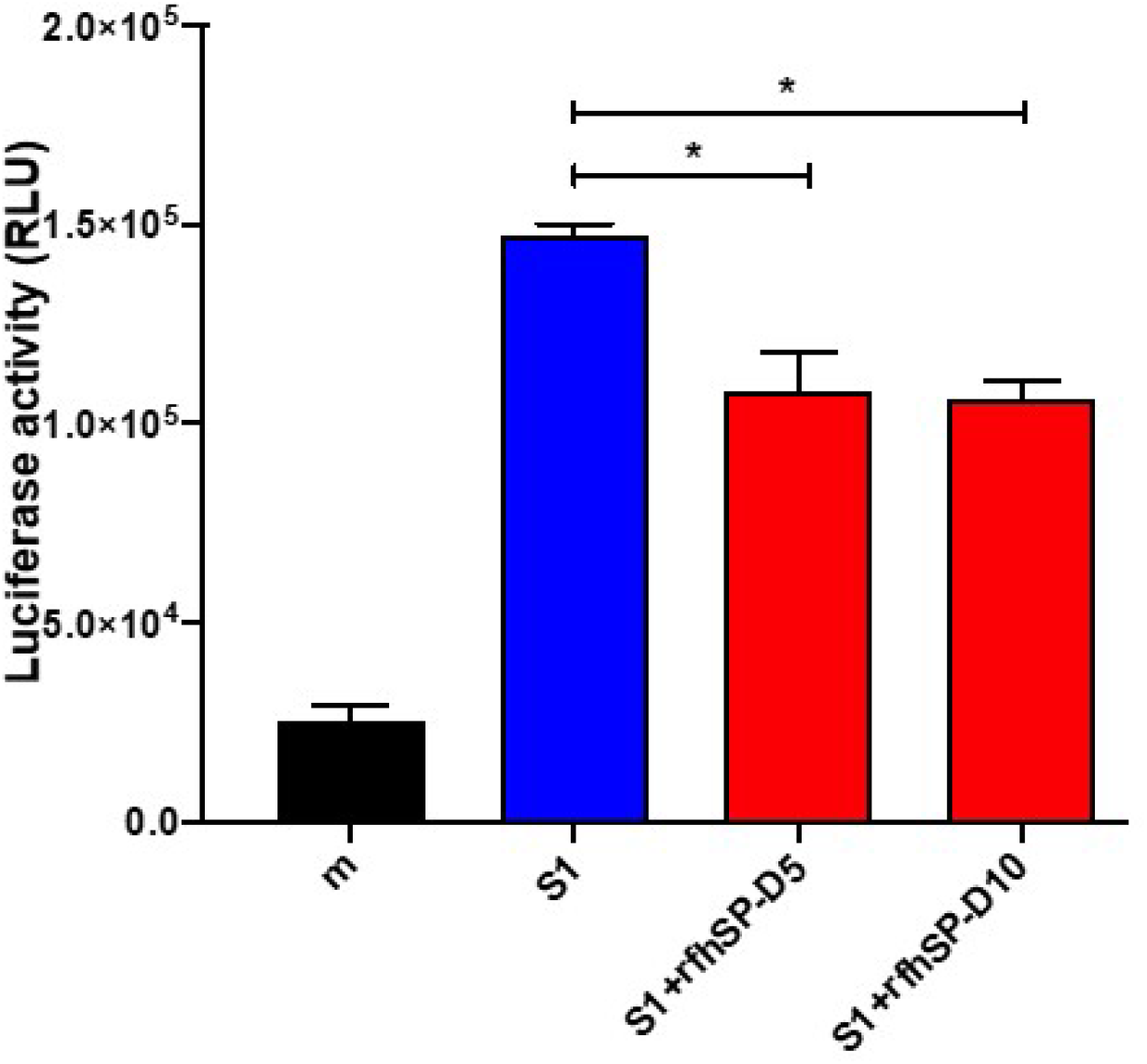
rfhSP-D act as an entry inhibitor of SARS-COV-2 infection. Luciferase reporter activity of rfhSP-D treated HEK293T cells (overexpressing ACE2 receptor) transduced with of SARS-CoV-2 S1 pseudotyped lentiviral particles. Significance was determined using the unpaired one-way ANOVA test statistical analysis. (*p < 0.05) (n = 3).

## Discussion

The innate immune system has evolved cellular and molecular defense mechanisms critical for the recognition and restriction of SARS-CoV-2-mediated respiratory tract infections, and for the activation of subsequent adaptive immune responses (32). SARS-CoV-2 infection is responsible for a higher transmissibility, mortality and morbidity rates that has caused the current global pandemic (33–35). Given that the SARS-CoV-2 is a newly emerged pandemic infection, it is fundamentally crucial to understand the roles of host immune response-innate as well as adaptive wings, which in turn, is likely to have profound impact on designing and developing effective anti-SARS-CoV-2 therapies.

The primary target for SARS-CoV-2 is the alveolar epithelial type II cells (36), and the viral entry to the host cell is mediated by the ACE2 receptor (37). Furthermore, viral entry into the host cells also depends on the activity of TMPRSS2 protease as it facilitates S protein cleavage into S1 and S2 portions. This enables S-mediated targeting and receptor-mediated early fusion pathway driven by the S2 subunit (19). Human SP-D is a lung collectin, synthesized by Clara cells (38) and alveolar type-II cells (1). Anti-viral role of SP-D has been reported against HIV-1 and IAV infection (5, 6, 8, 39, 40). In addition, increased serum SP-D has been observed in SARS-CoV patients (11); interaction between SP-D and the S protein of SARS-CoV leads to enhanced phagocytosis (11, 12). Furthermore, SP-D can also inhibit viral infection of 16HBE cells infected with HCoV-229E (13). However, SP-D-mediated inhibition of SARS-CoV infection and its subsequent immune response is not fully studied. Therefore, this study was aimed at examining the ability of rfhSP-D to act as an entry inhibitor of SARS-CoV-2 infection using pseudotyped lentiviral particles expressing SARS-CoV-2 S1 protein. Being a potent innate immune molecule present in the lung surfactant, SP-D is expected to play an important protective role in the pathogenesis of COVID-19.

In this study, affinity purified, and LPS-free rfhSP-D was found to interact with S1 protein and its RBD of SARS-CoV-2 in a dose-dependent manner. SARS-CoV-2 However, the interaction between rfhSP-D and RBD was stronger compared to that with S1. Inhibition of rfhSP-D binding to S protein by EDTA or maltose suggested that rfhSP-D bound to the carbohydrate moieties on S protein of SARS-CoV-2 (12). We also examined whether rfhSP-D treatment can inhibit the interaction of SARS-CoV-2 S1 with ACE2 receptor on HEK293T cells. SARS-CoV-2 S1 protein (2 μg/ml), pre-incubated with a varied concentration of rfhSP-D (0.625 −10 μg/ml), showed reduced binding to HEK293T cells overexpressing ACE2 receptor in a dose-dependent manner.

Targeting viral entry into a host cell is an emerging approach for designing and developing anti-viral therapies as viral propagation can be either restricted or blocked at an early stage of viral cycle, diminishing drug resistance by released viral particles. In this study, we examined the entry inhibitor role of rfhSP-D against SARS-CoV-2 by luciferase reporter assay. Pseudotyped lentiviral particles were generated as a safe alternative method to mimic the structural surface of SARS-CoV-2, and to test whether rfhSP-D treatment can promote or prevent viral entry into the host cells. Approximately 0.5 RLU fold reduction was seen with rfhSP-D (5 or 10 μg/ml) treatment when compared to untreated sample (1 RLU fold; Cells + SARS-CoV-2). A significantly reduced luminescent signal following rfhSP-D treatment significantly indicates that the interaction of rfhSP-D with SARS-CoV-2-S1 and its putative receptor ACE2 restricted the binding and entry of the virus, suggestive of an entry inhibitory role of rfhSP-D against SARS-CoV-2 infection.

SARS-CoV-2-mediated lung injury is correlated with diffuse alveolar damage and air space oedema, thus, accompanied by interstitial infiltration of inflammatory cells, trigger of coagulation, and fibrin deposition (41–44). Potential biomarkers to be considered during SARS infection include increased levels of inflammatory plasma makers, coagulation, and fibrinolysis (45, 46). Damage to the alveolar epithelial barrier is a characteristic feature of an acute respiratory distress syndrome (ARDS) and acute lung injury (ALI); levels of plasma surfactant proteins such as SP-A and SP-D may have a prognostic value (47–49). Thus, this study prompts further investigation into the role of pulmonary surfactant in COVID-19.

In summary, rfhSP-D, containing homotrimeric neck and CRD regions, acts as an entry inhibitor of SARS-CoV-2 infection by restricting the viral entry into HEK293T cells overexpressing ACE2 receptor. Time is ripe for taking the knowledge about the involvement of rfhSP-D and its associated anti-viral effects forward to the development as a novel therapeutic approach to target multiple cellular signaling pathways. The mechanisms which enable rfhSP-D to trigger anti-viral effect are viral specific due to the differential effects and variation on the cell types and presence of putative receptors. There is a clear therapeutic potential of rfhSP-D against SARS-CoV-2 where increased glycosylation leads to evasion of antibody susceptibility, but increased susceptibility against soluble pattern recognition receptors (PRRs) such as SP-D. Having established the specific nature of interactions between rfhSP-D and SARS-CoV-2, we hope to examine host response in the murine models of infection using wild type and SP-D knock-out mice in future.

